# DDR and TMEM132 genes control progenitor targeting and regenerative assembly of the planarian eye

**DOI:** 10.64898/2026.10.03.756328

**Authors:** Chaeli J. Place, Katherine C. Lo, Eleanor G. Clark, Christian P. Petersen

## Abstract

How progenitors can faithfully self-assemble to build new organs in adulthood is a central unsolved problem in regeneration biology. The planarian eye is a model for uncovering principles of *de novo* organogenesis during regeneration. In this process, pluripotent adult stem cells specify into *ovo*+ migratory progenitors which target to the eye field due to Wnt and FGFRL signals produced from bodywall muscle, then differentiate into photoreceptor neurons (PRNs) and pigmented optic cup (OC) cells, followed by organization of these two cell types in an opposing pattern that completes eye organ maturation. We used scRNA-seq to characterize eye progenitor transcriptomes during regeneration, followed by FISH validation, and RNAi screening to identify regulators of eye regeneration. This uncovered a function for the discoidin domain receptor *ddr-1* in preventing the premature terminal differentiation of migratory eye progenitors. DDRs encode collagen receptors, and bodywall muscle is the major source of collagen in planarians, suggesting eye progenitors likely interface with muscle ECM for their proper deployment in regeneration. We further identified a novel eye morphogenesis phenotype from knockdown of *tmem132-1*, causing failed heterotypic adhesion between eye cell types. TMEM132 factors are conserved cell-surface receptors that mediate adhesion and regulate migration, and human TMEM132 mutations are associated with several neurobehavioral diseases. Therefore, planarian *tmem132-1* likely directly regulates the capture and assembly of migratory progenitors into mature eyes during regeneration. Together, these results define new steps in the eye regeneration pathway and suggest ancient roles for DDR and TMEM132 in progenitor migration and coordinated organ assembly.

## INTRODUCTION

Regeneration must not only generate new differentiated cells but also assemble them into functional tissues and organs. Planarians, which can regenerate any missing tissues following surgical injuries, represent an extreme example of this ability (Reddien, 2018). In these organisms, an abundant pool of adult pluripotent stem cells called neoblasts is broadly distributed in the animal and differentiates into all adult cell lineages through the production of lineage-specified subpopulations. Progenitors for distinct lineages are spatially intermingled in planarians (Park et al., 2023), so regeneration likely involves a substantial role for targeting many distinct precursor cells to their appropriate locations followed by robust assembly processes to reproducibly form mature tissues. A significant gap persists in understanding these molecular processes enabling *de novo* organ assembly in a post-embryonic context.

The planarian eye serves as a powerful model to dissect the mechanisms of adult organ regeneration (Reddien, 2021). Throughout their lifespan, planarians regenerate their eyes through continuous homeostatic turnover and rebuild eyes *de novo* following injury. The planarian eye is primarily composed of two cell types, *tyosinase+* pigmented optic cup (OCs) cells that organize to form a medial hemisphere, and lateral photoreceptor neurons (PRNs) which project axons ventrally into the brain for transducing light-dependent behavior. Dendritic projections from photoreceptors localize dorsally and coincide with an unpigmented transparent region, which is thought to facilitate light transmission (Scimone et al., 2026). Eye regeneration and homeostatic maintenance require a common population of migratory post-mitotic eye progenitors expressing transcription factors *ovo, sine oculis* (*six-1/2*) and *eyes absent* (*eya*) (Lapan and Reddien, 2011; Lapan and Reddien, 2012) which derive from the Piwi+ neoblast pluripotent adult stem cells that produce all tissue lineages in adult planarians (Wagner et al., 2011). PRN differentiation requires the transcription factor *otxA*, while OC cell differentiation requires *sp6-9* and *dlx,* and these factors are expressed in subpopulations of migrating *ovo+* progenitors (Lapan & Reddien, 2011). Additional regulators include *p66,* which suppresses PRN differentiation (Vasquez-Doorman and Petersen, 2016); factors participating in terminal differentiation of OC cells (*inx-14, melanoir*, *egfr-1*) and PRNs (*klf*, *rgs-1*, *gcr591*, *FoxQD, soxB-1, smad6/7-2*) (Gonzalez-Sastre et al., 2012; Lapan and Reddien, 2012; Scimone et al., 2026); and factors that promote the PRN rhabdomere projections between OCs and PRNs (*ft-1, ablim, ddr-4, ankyrin-2)* (Scimone et al., 2026).

A complex patterning system governs the spatial control of eye differentiation through body-wall muscle signaling. Wnt and FGFRL factors are expressed in domains across the anteroposterior axis (Witchley et al., 2013) where they regulate body regionalization, including the location of new eye regeneration in the anterior. Inhibition of *wnt11-6/wntA* or the FGF receptor-like factor *nou darake (ndk)* leads to posterior ectopic eyes, while inhibition of the Wnt antagonist *notum* produces anterior ectopic eyes and anterior brain tissue (Cebria et al., 2002; Adell et al., 2009; Hill and Petersen, 2015; Scimone et al., 2016; Hill and Petersen, 2018). A separate system of trunk patterning genes (*wntP-2, ndl-3*, and *ptk7*) is responsible for patterning of the pharynx (Lander and Petersen, 2016; Scimone et al., 2016). Src signaling inhibition causes posteriorization of both the eyes and the pharynx, indicating Src likely acts through a common parallel or downstream pathway for both head and trunk patterning (Bonar et al., 2022). Perturbation of muscle differentiation itself can generate eye placement defects, supportive of the upstream role this tissue plays in positional information for regeneration (Scimone et al., 2017; Scimone et al., 2018; Scimone et al., 2020).

A separate set of processes likely governs the ability for progenitors to migrate, hone to proper destinations, and be captured by eyes. Generating eyes in incorrect locations either through transplantation or shifts to the patterning system can allow sustained eye homeostasis at locations outside of the target zone, where the regeneration system produces new eyes (Atabay et al., 2018; Hill and Petersen, 2018). In addition, eye resections followed by counting eye cells showed that nearby eyes can compete with each other for the acquisition of progenitors (Hill and Petersen, 2018). These observations suggest that upstream patterning systems used during regeneration are likely distinct from the ability of organs to capture progenitors and that differentiation may occur at the cessation of the migratory state. *map3k1* is required to prevent premature terminal differentiation of migratory progenitors, and its inhibition results in ectopic posterior eye formation in the region transited by progenitors, likely downstream of the muscle-based patterning system (Lo and Petersen, 2025; Canales et al., 2026). Likewise, integrin inhibition causes chaotic differentiation of eyes, neurons, and other tissue in incorrect locations, suggesting a role for extracellular matrix (ECM) interactions for proper cell targeting (Bonar and Petersen, 2017; Seebeck et al., 2017). However, the upstream and downstream factors sustaining progenitor migration or involved in eye cell chemotaxis to target locations are unknown. The progenitor capture process has additionally been studied by resecting eyes in regenerating head fragments, which transiently shift their eye target zone as they rescale to regenerate into a smaller body (Atabay et al., 2018). Inhibition of several adhesion factors (*pcdh-5*, *BOC*, *ft-1*, *pcdh-6*, *pcdh-7*, *inx-4*, and *TMEM258*) induces the formation of ectopic anterior eyes only following head amputation and removal of a pre-existing eye, likely because stray progenitors fail to be captured and initiate an ectopic eye anteriorly under these conditions (Scimone et al., 2026). Intriguingly, no factor has yet been defined whose inhibition completely prevents either homotypic interactions within each eye cell type or heterotypic interactions potentially involved in associating the distinct cell types together. Proper morphogenesis of photoreceptors and OCs is mutually dependent (Lapan and Reddien, 2011), and whole eyes can be nucleated either by transplantation of PRNs or OC cells (Scimone et al., 2026). Therefore, cross-interactions of eye cell types are likely critical for eye morphogenesis, although these remain unknown.

We conducted RNA interference (RNAi) screening of eye-expressed genes defined by single-cell RNA sequencing (scRNAseq) and identified novel roles for two genes. We confirm prior analysis that inhibition of a collagen receptor (discoidin domain receptor 1, *ddr-1*) causes loss of the transparent region around the eyes and generates ectopic anterior eyes as part of its role in progenitor capture. We found that inhibition of this gene leads to ectopic posterior eyes due to premature differentiation of migrating progenitors and confirmed that *ddr-1* is expressed in *ovo+* progenitors, suggesting these cells might directly use collagen-rich ECM contact to sustain migration ability. We also identified an adhesion receptor *tmem132-1* required for the association of PRNs and OC cells and essential for building the eye. These results define steps in the process of regenerative morphogenesis.

## RESULTS

### RNAi screen to identify regulators of eye assembly

To identify genes with potential roles in progenitor migration and eye assembly, we surveyed the transcriptomes of eye progenitors and mature eye cells using scRNAseq. We chose to profile two timepoints of head regeneration (5 and 8 days post amputation [dpa]) known to be enriched for the presence of migratory *ovo+* cells and the active assembly of new eyes (Figure 1A) (Lapan & Reddien, 2012). In total, 60,816 cells were profiled (Figure 1B) and subclustered by expression of either *ovo* or *six-1/2* resolved clusters of *piwi-1+* eye-specified neoblasts, *otxA+* post-mitotic progenitors, *opsin+* PRNs, and *tyrosinase+* OC cells (Figure 1C). *sp6-9+piwi-1+* cells represented a subset of eye-specified neoblasts, although no post-mitotic *sp6-9+* progenitors were identified, suggesting this population may differentiate directly. We performed differential gene expression analysis with the goal of identifying factors enriched in eye progenitors. We investigated 61 genes, focusing on cell-surface receptors, adhesion molecules, transcription factors, and other signaling molecules, and verified expression of 36 factors using double fluorescence in situ hybridization (FISH) with *ovo* in 5 dpa regenerating trunk fragments in which progenitors and newly differentiated eye cells are present (Figure 1D). Similar to prior analyses of the planarian eye and other differentiation systems in planarians (Lapan and Reddien, 2011; Lapan and Reddien, 2012; Scimone et al., 2014), the FISH and scRNAseq data found that all genes expressed in differentiating eye progenitors also had expression in mature eye cells (Figure 1D, Figure S1).

**Figure 1.**
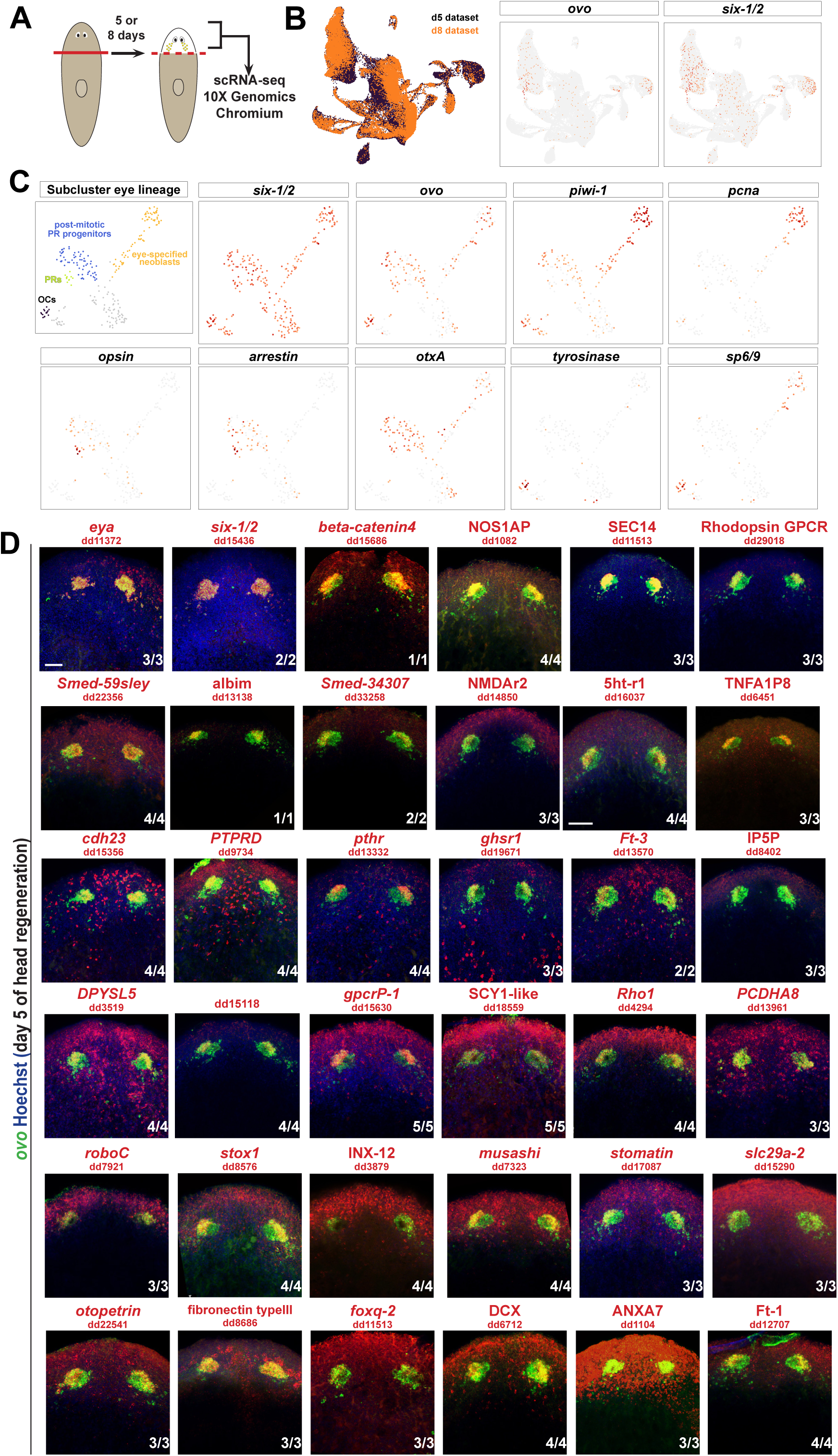
Single-cell RNAseq and FISH to identify genes expressed in the planarian visual system. (A) Surgical strategy to perform scRNAseq on regenerating head blastemas to recover eye progenitors (yellow). (B) Left, UMAP depicting profiled cells. Right, *ovo* and *six-1/2*-expressing cells. (C) Subclustering *ovo+/six-1/2+* cells to identify eye specified *piwi-1*(high) proliferative (*pcna*+) neoblasts (orange), *piwi-1*(low) post-mitotic PRN progenitors expressing *otxA* (blue), differentiated PRNs expressing *opsin* and *arrestin* (yellow), and differentiated OC cells expressing *tyrosinase* (black) (and unassignable cells marked in gray). While we recovered *ovo+piwi1+sp6-9+* neoblasts specified toward OC fates, we did not recover any post-mitotic *sp6-9+* OC progenitors, suggesting the rarity of this hypothesized cell population. (D) Double FISH to detect expression of identified factors along with *ovo* in day 5 regenerating trunk fragments undergoing eye regeneration. Scorings indicate the number of animals appearing as displayed. Scale bar, 50 microns.

To test whether any candidates identified through scRNAseq regulate eye regeneration, we perturbed their expression through RNAi. Animals were fed liver combined with double-stranded RNA (dsRNA) six times over two weeks then amputated into head, trunk, and tail fragments to observe regeneration phenotypes. Live animals were scored for eye regeneration defects and fixed at 9 dpa. Known regulators of eye regeneration, including *eya*, *six-1/2*, and *robo-C*, and *meis* showed eye regeneration defects in accordance with their published roles (Table S2)(Lapan and Reddien, 2011; Lapan and Reddien, 2012; Scimone et al., 2020). Of the 61 genes screened, we identified four genes with unpublished roles in planarian eye regeneration whose knockdown produced a variety of eye regeneration defects (Table S1). *ddr-1* RNAi and *tmem132* RNAi produced the most severe, novel defects in eye morphology and assembly and were thus prioritized for subsequent analysis.

### *ddr-1* is required for planarian eye regeneration and maintenance

Discoidin domain receptors (DDRs) are receptor tyrosine kinases activated by interactions with collagen in order to regulate cell-ECM interactions (Leitinger, 2014). Knockdown of *ddr-1* led to the loss of the transparent region, leading to a “whiteless” eye phenotype across fragment types, and the production of ectopic eyes in head fragments (5/9, 55%) and more weakly in trunk fragments (1/9, 11%) (Figure 2A). While the transparent region loss and formation of anterior ectopic eyes are consistent with prior findings (Scimone et al., 2026), the occurrence of posterior ectopic eyes is a newly reported phenotype. Whereas anterior eyes can indicate an anterior shift to patterning systems similar to *notum(RNAi)* phenotypes, prior analysis of *ddr-1* RNAi found that eye removal in these animals did not cause a shift to the location of regeneration and instead *ddr-1* likely has a previously determined role in progenitor capture (Scimone et al., 2026). We were intrigued by posterior eye formation, which can occur if progenitors differentiate prematurely, similar to *map3k1* RNAi (Lo and Petersen, 2025; Canales et al., 2026). First, we tested whether *ddr-1* executed this function only in regeneration or also in homeostatic conditions. Long-term knockdown of *ddr-1* in uninjured animals also caused the loss of the transparent region by 3 weeks (8/8 animals) and ectopic anterior and posterior eyes (4/8 animals) (Figure 2B). By 5 weeks, all animals had at least one ectopic eye (8/8 animals), and several had multiple ectopic eyes. Thus, *ddr-1* also regulates the location of eye formation in homeostatic contexts. To examine the location of ectopic eye regeneration in these animals systematically, we determined the relative locations of ectopic eyes across animals of different sizes by normalizing anteroposterior and mediolateral positions to the original eyes in head fragments and homeostatic animals (Figure 2C). This analysis confirmed that *ddr-1* RNAi causes eyes to form at both anterior and posterior positions with respect to the original eyes, and the similarity of this process in head fragments versus uninjured animals indicates these properties are not dependent on the regeneration-induced axis rescaling occurring during morphallaxis.

**Figure 2.**
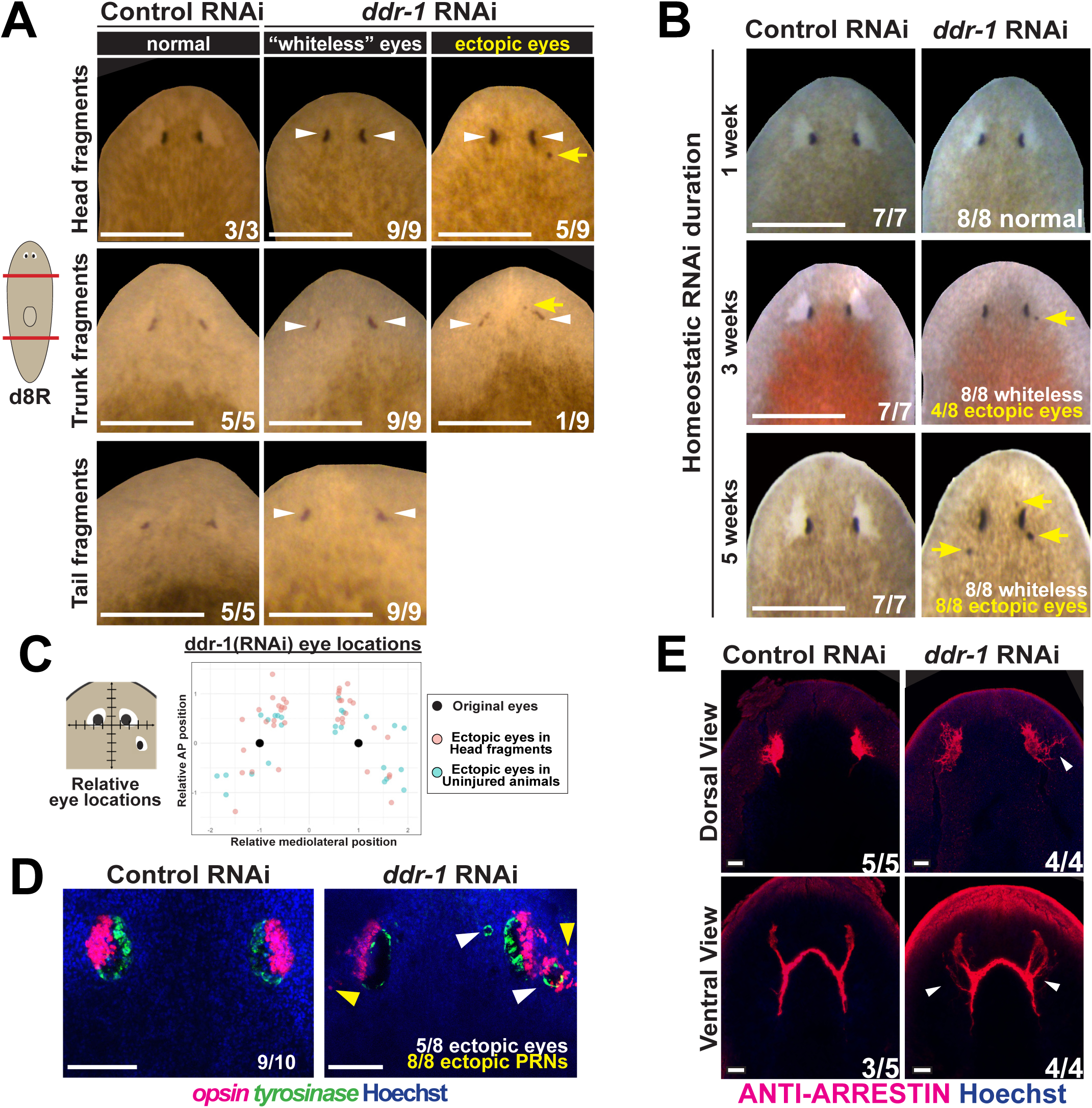
*ddr-1* RNAi causes formation of ectopic posterior and anterior eyes. Inhibition of *ddr-1* in regenerating (A) or uninjured (B) animals caused loss of the transparent region (white arrowheads) and formation of ectopic posterior and anterior eyes (yellow arrows). (C) Analysis of ectopic eye positions in *ddr-1(RNAi)* animals with respect to the pre-existing eyes (black dots) normalized by animal size. In homeostatic animals or regenerating head fragments, ectopic eyes formed anteriorly and/or posteriorly. (D) *ddr-1(RNAi)* animals stained by FISH detecting ectopic eyes containing *opsin+* PRNs and *tyrosinase+* OC cells (white arrowheads), as well as isolated misplaced *opsin+* cells (yellow arrowheads). (E) Anti-arrestin staining detected ectopic PRN axons projecting toward the brain and disorganized dorsal dendrites. Scorings indicate the number of animals appearing as displayed. Scale bars, 500 microns (A-B), 100 microns (D), 50 microns (E)

To gain understanding of the nature of this phenotype, we further examined *ddr-1(RNAi)* animals for their PRNs and OC cells using FISH and immunostaining with VC-1 antibody to detect ARRESTIN (Figure 2D-E). Ectopic eyes from *ddr-1(RNAi)* animals contained both PRNs and OC cells and projected ectopic axons toward the brain (Figure 2D-E). *ddr-1(RNAi)* animals also contained stray PRNs (i.e., *opsin+* cells) not associated with complete eyes, potentially indicating a lack of spatial control over PRN differentiation. These ectopic PRNs were accompanied by ectopic PRN axons projecting toward the brain. Confirming prior analysis (Scimone et al., 2026), the dorsal field of photoreceptor dendrites normally associated with the transparent region were also disorganized (Figure 2E). Together, *ddr-1* RNAi results in the formation of mature eye cell types at inappropriate locations.

### *ddr-1* suppresses the terminal differentiation of photoreceptor neuron progenitors

The analysis above suggested a possible model in which progenitors use DDR-1 during the migration process to prevent premature terminal differentiation, in addition to its roles in forming the transparent region and allowing for the capture of migratory progenitors into mature eyes. To examine this hypothesis, we first use FISH to validate the measurements from scRNAseq showing that *ddr-1* is expressed in *ovo+* progenitors during regeneration. In animals fixed in the process of forming eyes at 5 days of head regeneration, *ddr-1* expression was detected both in the maturing eyes as well as in *ovo+* cells in neighboring trails of migratory eye progenitors (Figure 3A). We next tested whether *ddr-1* RNAi impacted the expression states within progenitors using double FISH to detect *opsin* and *ovo* on day 5 of head regeneration. Double-positive *opsin*+*ovo*+ cells in the progenitor trails were significantly detected in *ddr-1(RNAi)* animals but not in control animals (Figure 3B). Collectively, this evidence reveals that *ddr-1* is expressed in migrating progenitors and is necessary to prevent premature terminal differentiation of these cells as they migrate toward their target destination.

**Figure 3.**
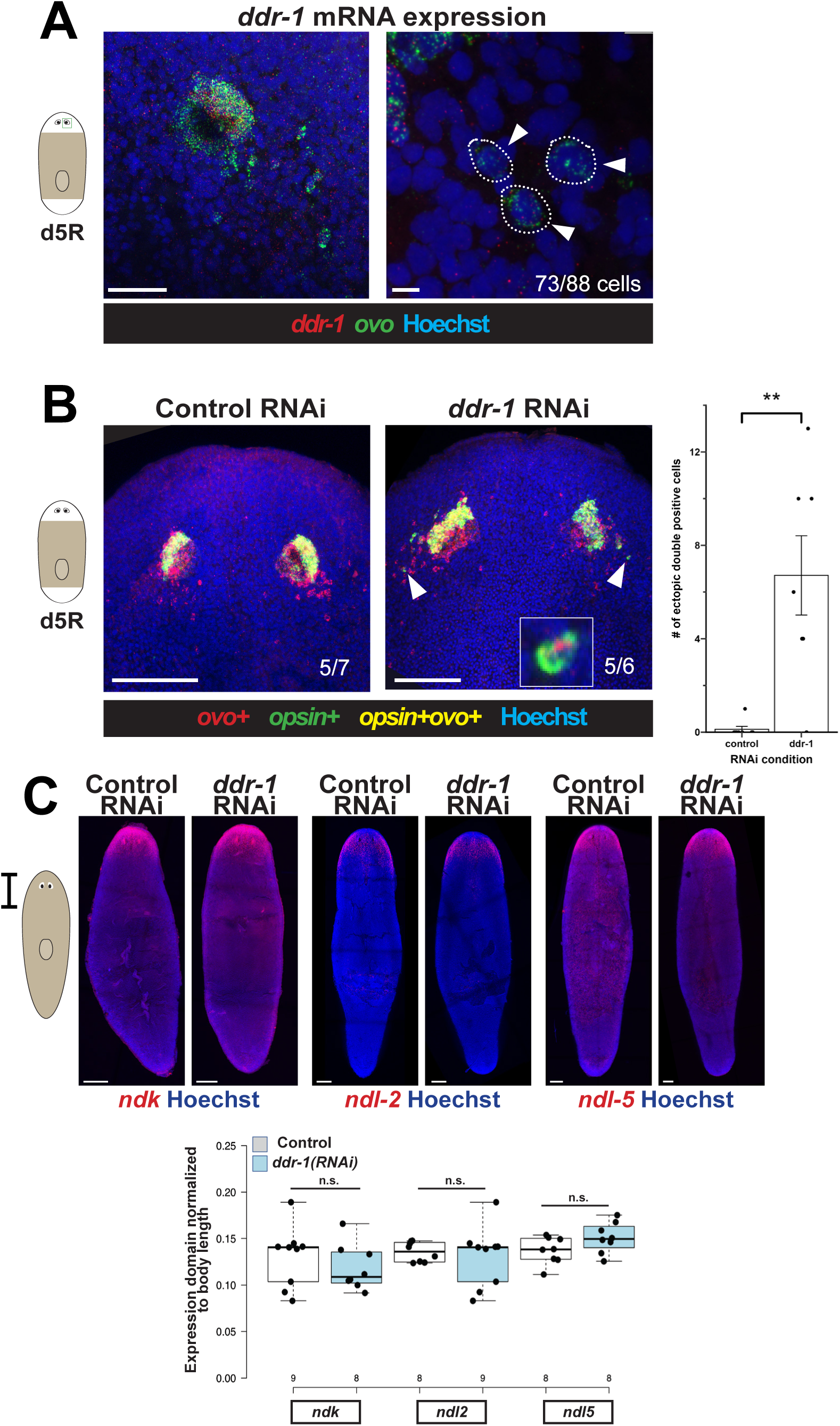
*ddr-1* is expressed in migratory eye progenitors and prevents premature terminal differentiation. (A) Double FISH detected expression of *ddr-1* within *ovo*+ eye progenitors detected in day 5 of head regeneration. Left panel shows regenerating eye imaged at 20x and right panel shows zoom-in of *ovo*+ cells located in the progenitor migratory zone imaged at 40x. 83% (73/88) of dispersed *ovo+* cells counted were *ddr-1+*. (B) Double FISH detecting that *ddr-1(RNAi)* animals have increased numbers of migratory *ovo+* that co-express *opsin* (white arrowheads, inset shows zoom-in image of a double positive cell), while such cells were very rarely identified in control animals. Right, quantification of *opsin+ovo+* cells outside of the mature eyes (**, p < 0.01 by two-tailed t-test). (C) FISH detecting anterior patterning factors expressed in regional domains within muscle (*ndk*, *ndl-2*, *ndl-5*). *ddr-1* RNAi animals did not have visible differences in the size and position of these domains. Bottom, quantification of the size of each anterior expression domain, normalized to body length also did not find significant differences in the expression of patterning factors from muscle after *ddr-1* RNAi. Statistical tests show two-tailed t-test, and numbers across the x-axis show sample sizes (n.s., p>0.05). Scorings indicate the number of animals appearing as displayed. Scale bars, 50 microns (A, left), 5 microns (A, right), 100 microns (B), 200 microns (C).

This data suggested a model in which *ddr-1* functions in eye regeneration to control differentiation downstream of the global patterning systems signaled from body-wall muscle. To evaluate this possibility, we examined the expression of muscle-expressed patterning factors marking domains within the anterior. *ddr-1* RNAi did not visibly influence expression domains of *ndk*, *ndl-2*, and *ndl-5*, and quantification of these domain sizes did not reveal any difference (Figure 3C). Consistent with these results, *ddr-1(RNAi)* animals did not have increased brain sizes as assessed by numbers of *cintillo+* neurons (Figure S2). By contrast, RNAi of head patterning factors *ndk* or *wntA/wnt11-6* cause ectopic posterior eyes along with brain expansion (Cebria et al., 2002; Kobayashi et al., 2007; Adell et al., 2009; Hill and Petersen, 2015; Scimone et al., 2016; Hill and Petersen, 2018). These results are consistent with a possible mechanism in which muscle provides signals dictating the placement of eyes in regeneration, while *ddr-1* acts at a downstream step within migrating progenitors to suppress terminal differentiation while they are in transit. DDRs are known to bind and be activated by collagen (Leitinger, 2014), so this mechanism may involve progenitor interactions with collagen to sustain the migratory state prior to reaching their destination.

### TMEM132 regulates the association of photoreceptor neurons and optic cup cells

Our screen identified another novel phenotype of specific disruption to the eye regeneration program from the inhibition of *tmem132-1*. The TMEM132 family is a highly conserved group of single-pass transmembrane proteins (Sanchez-Pulido & Ponting, 2018) (Figure S3A). Both our scRNAseq analysis (Figure S1) and analysis from a prior scRNAseq dataset (Scimone et al., 2026) (Figure S3B) indicated that *tmem132-1* is expressed in post-mitotic progenitors and PRNs but only weakly in OC cells. Consistent with these data, using FISH we found that *tmem132-1* was expressed in the majority of *ovo*+ eye progenitors (80/96 cells) and laterally at the location of PRNs within the eye (Figure 4A).

**Figure 4.**
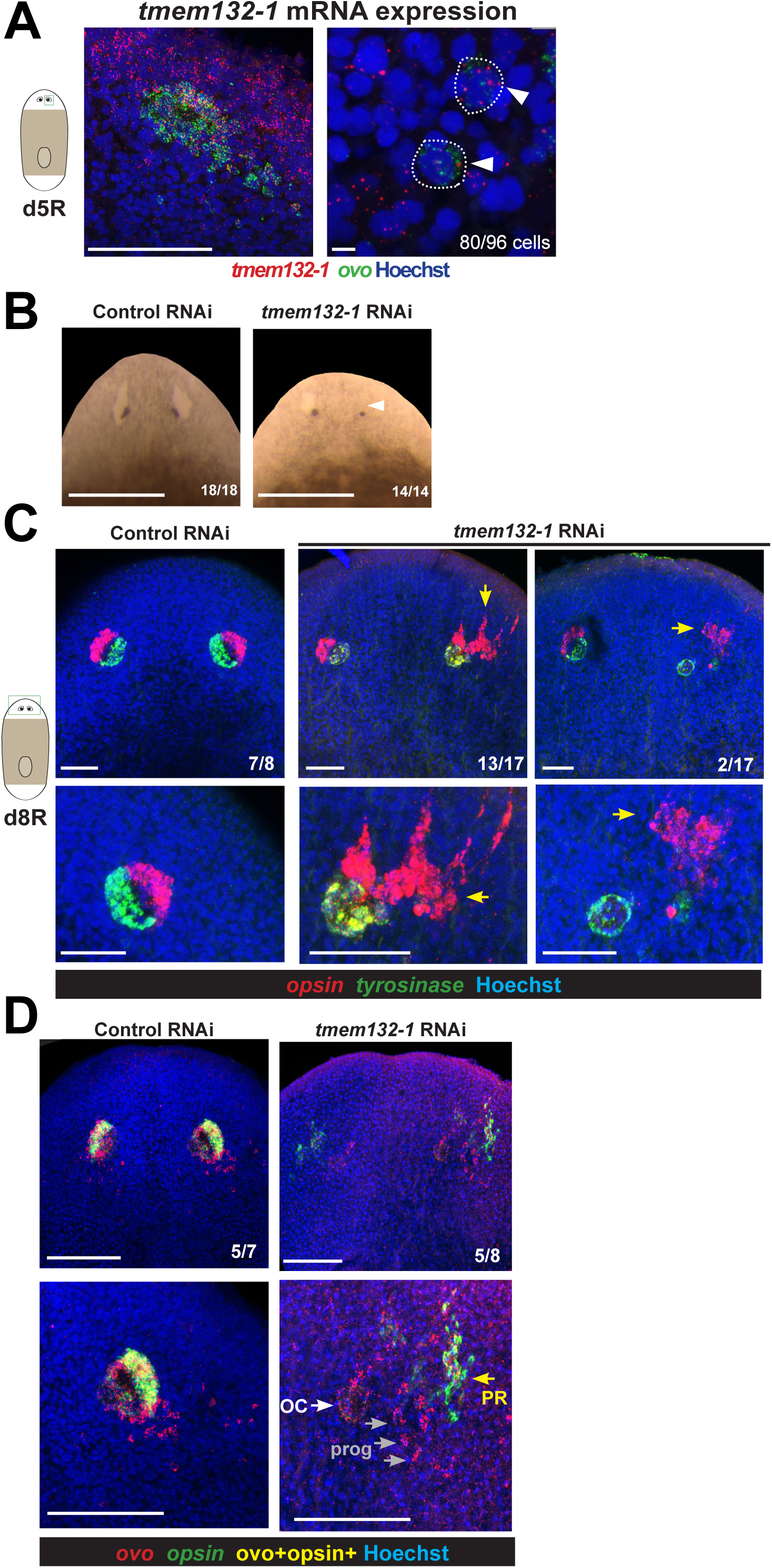
*tmem132-1* regulates the assembly of eye cell types. (A) FISH detecting expression of *tmem132-1* in *ovo+* eye cells (left, images taken at 20x) and also in *ovo*+ migratory eye progenitors (right, images taken at 40x) imaged in animals at day 5 of head regeneration. 83% (80/96) of *ovo*+ progenitors expressed *tmem132-1*. (B) *tmem132-1* RNAi caused defects in the ability to form the transparent region of the eye (white arrowhead) without affecting the pigmentation of OCs. (C) Double FISH showing morphogenesis defects in the assembly of eyes after *tmem132-1* RNAi. *tyrosinase+* OC cells formed enclosures while *opsin+* PRNs aggregated away from OCs (middle panels, 13/17 animals) and in some cases were not connected to OCs at all (right panels, 2/17 animals) (yellow arrows). (D) Dispersed *ovo+* cells likely representing eye progenitors (gray arrows) were located between the circular OC (white arrow) and the detached PRNs (yellow arrow) in day 5 *tmem132-1(RNAi)* trunk fragments regenerating new eyes. Scorings indicate the number of animals appearing as displayed. Scale bars, 50 microns (A, left), 5 microns (A, right), 500 microns (B), 100 microns (C), or 50 microns (D).

*tmem132-1* RNAi caused a novel phenotype of dysregulated eye morphogenesis. In live animals, the transparent region was reduced or absent (14/14 animals) (Figure 4B). About 36% of the time, animals had an asymmetric appearance with respect to the amount of transparent region lost. In live animals, pigmented OCs were present at apparently normal locations. FISH staining of *tmem132-1(RNAi)* animals revealed that *opsin+* PRNs accumulated away from the *tyosinase+* OC cells (13/17 animals) and in some cases were entirely detached from them (2/17 animals) (Figure 4C). *tyrosinase+* cells additionally tended to form an enclosed ball rather than their typical hemisphere morphology (11/17 animals). Notably, both types of cells still appeared capable of associating with like types after *tmem132-1* RNAi, because *opsin+* cells localized near each other rather than dispersing spatially (Figure 4C). Therefore, *tmem132-1* is required for heterotypic association between PRNs and OC cells but likely not for the homotypic association within each cell type.

Similar phenotypes of rounded OCs have been observed after elimination of PRNs through *otxA* RNAi (Scimone et al., 2026). Likewise, phenotypes of mislocalized PRNs arise after elimination of OC cells through *sp6-9* RNAi (Scimone et al., 2026). *tmem132-1* RNAi causes similar defects in OC and PR morphogenesis without the loss of either cell type, indicating *tmem132* has a role in allowing the two cell types to associate without controlling their formation. This specific defect in the process of morphogenesis appears to be rare, because RNAi screening of over 450 eye-expressed factors and other prior RNAi screening in the literature did not previously identify such a phenotype (Reddien et al., 2005; Lapan and Reddien, 2012; Scimone et al., 2026). Progenitors in *tmem132-1(RNAi)* appeared to distribute to the region between OCs and the detached PRNs (Figure 4D), suggesting the intriguing possibility that each differentiated cell type might individually be capable of attracting their own progenitors. Based on the rarity and specificity of the *tmem132-1* phenotype, as well as the context *tmem132-1’s* molecular function as an adhesion receptor, we posit that the interaction between PRNs and OC cells could be mediated directly by the TMEM132-1 protein.

## DISCUSSION

The observations described here suggest two candidate models accounting for the action of *ddr-1* and *tmem132-1* in regenerative eye assembly. The *ddr-1* RNAi phenotype is distinct from other ectopic eye phenotypes produced by knockdown of position control genes, such as *notum, ndk, wntA/wnt11-6*, or *src-1* (Cebria et al., 2002; Adell et al., 2009; Hill and Petersen, 2015; Scimone et al., 2016; Hill and Petersen, 2018; Bonar et al., 2022). First, in *ddr-1* RNAi, ectopic eyes do not form only either anterior or posterior to existing eyes, as would be expected if positional information were altered such that the target zone for eye regeneration shifts in one direction. Further, unlike other upstream regulators of head patterning, *ddr-1* appears to regulate the eye differentiation process rather than head regionalization in general. Finally, the *ddr-1* RNAi phenotype involves both the production of whole ectopic eyes and aberrant differentiation of isolated eye cells. Therefore, we suggest that *ddr-1* has multifunctional roles in the eye differentiation process. Normal eye regeneration involves eye nucleation in particular locations due to the influence of the body-wide patterning system, followed by capture of migratory progenitors to grow the eyes. In *ddr-1* RNAi, progenitors have a reduced ability to remain undifferentiated while they are in transit, and the production of new differentiated cells nucleates ectopic posterior eyes (Figure 5A). Therefore, *ddr-1* inhibition causes a phenotype that is partially overlapping with inhibition of *map3k-1*, in which both conditions lead to ectopic posterior eyes and premature differentiation of *ovo+* progenitors (Lo and Petersen, 2025). However, *ddr-1* likely has additional functions in eye regeneration, including the ability of existing eyes to capture progenitors in transit, so that its inhibition also leads to formation of ectopic anterior eyes and an inability for the PRNs to generate the transparent region of the eyes, as shown previously (Scimone et al., 2026). The molecular function of DDRs to bind collagen, along with the expression of *ddr-1* in *ovo+* trail cells, suggests that progenitors might sustain the migratory state by directly detecting the collagen-rich ECM (Figure 5B). Intriguingly, muscle is the primary source of collagen and most other ECM in planarians (Cote et al., 2019), suggesting that muscle might influence the differentiation process both through expressing body patterning factors like Wnt and FGF, and generating the template for progenitor transit.

**Figure 5.**
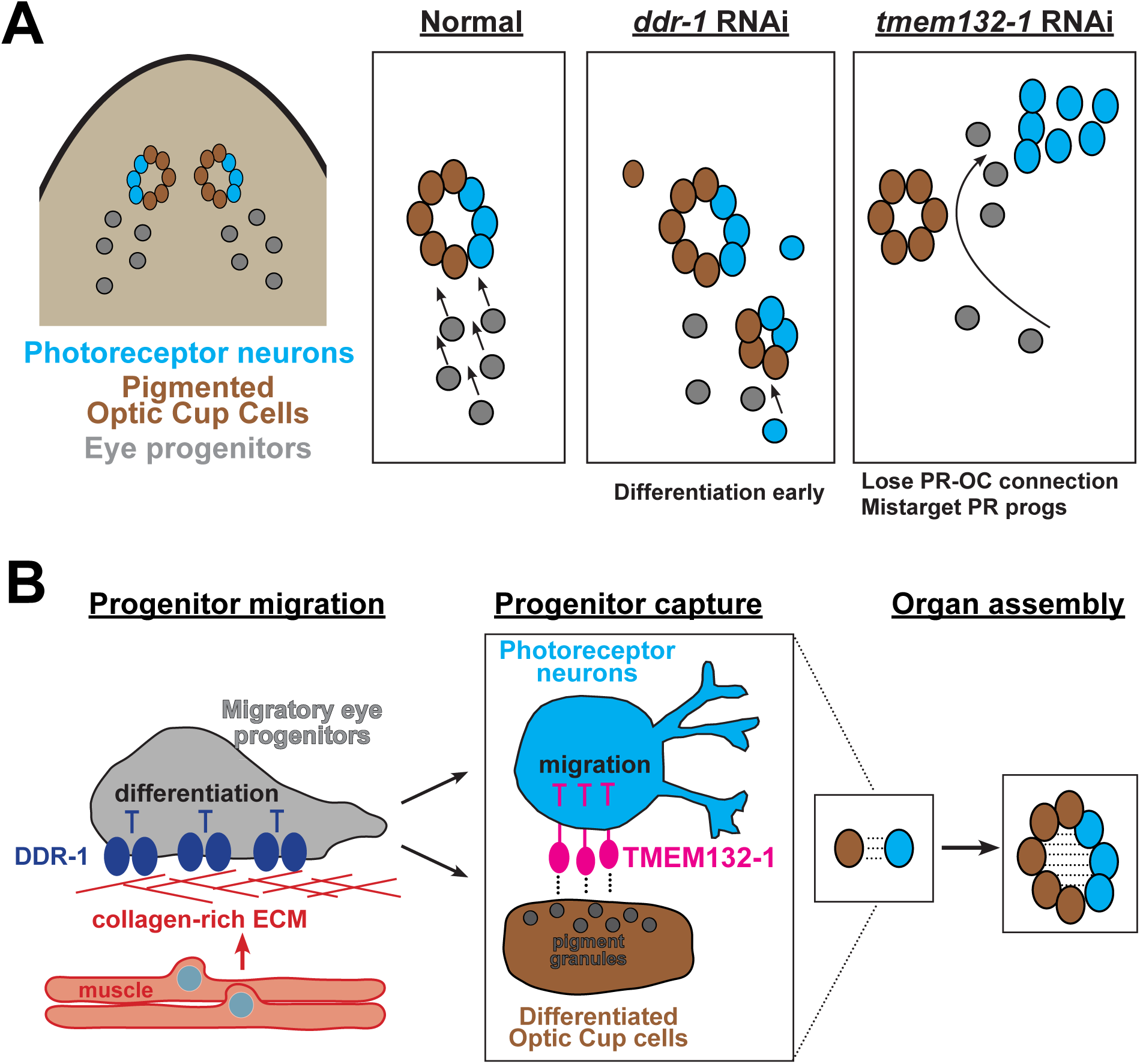
Model. (A) Diagram showing normal migration of eye progenitors and differentiation into pigmented OCs (brown) and PRNs (blue). *ddr-1* RNAi causes early differentiation of migratory progenitors and subsequent eye formation at ectopic posterior locations (shown), as well as defects for existing eyes to capture progenitors which continue to migrate and differentiate new eye cells at anterior locations. *tmem132-1* RNAi prevents the association between OC cells and PRNs while retaining the associations within each cell type, leading to defects in eye assembly. (B) A candidate cellular pathway for eye assembly. The expression of the collagen receptor *ddr-1* in migratory progenitors could enable these cells to sense connection to the collagen-rich ECM produced by muscle cells in order to suppress differentiation of these cells while in transit. Upon reaching a final destination dictated by Wnt/FGF signals from muscle in new eye regeneration (not shown), *tmem132-1* expressed from migratory eye progenitors as they mature into PRNs may help cease their migration by association with OC cells. Right, connections between eye cell types are critical for building the eye.

We furthermore define a novel step in the eye regeneration pathway regulated by *tmem132-1* to necessary for adjoining distinct cell types of the eye. The *tmem132-1(RNAi)* phenotype involves PRNs losing contact with OC cells, while homotypic PRN-PRN interactions and OC-OC cell interactions probably remain intact (Figure 5A). The expression of *tmem132-1* predominantly in progenitors and PRNs, along with its molecular function as an adhesion receptor, suggest it may act on these cells to directly enable association with OC cells through an unknown binding partner (Figure 5B). Alternatively, it is possible *tmem132-1* only indirectly allows for PRNs to be competent to adhere to OC cells, or that *tmem132-1* inhibition somehow unmasks other cell-cell adhesion modules that cause PRNs to adhere aberrantly to other nearby cells. Future work to identify possible TMEM132-1 binding partners and their cell type-specific expression patterns could help resolve these models. Additionally, the *tmem132-1(RNAi)* phenotype may provide more information about the process of eye nucleation downstream of the global patterning system. Although ectopically transplanted OC cells or PRNs can recruit the other cell type to form eyes (Scimone et al., 2026), the *tmem132-1(RNAi)* phenotype indicates that proper positioning of OC cells can take place without the correct localization of PRNs. Therefore, in normal eye regeneration, OC cells could be the initiating trigger for eye nucleation, followed by *tmem132-1*-dependent recruitment of PRNs. This model is also supported by the observation that *otxA(RNAi)* animals lacking PRNs place their OC cells in the appropriate locations, while PRNs appear somewhat misplaced anteriorly and laterally in *sp6-9(RNAi)* animals lacking OCs (Lapan and Reddien, 2011; Scimone et al., 2026). PRNs also engage in Notch signaling to trigger nearby progenitors to differentiate into glial cells (Scimone et al., 2025), so this model would imply that a typical eye assembly pathway involves the sequential placement of OC cells, recruitment of PRNs via *tmem132-1*, and finally PRN-dependent differentiation of eye glia cells.

Our findings also advance the understanding of TMEM132 factors in morphogenesis across organisms. Tmem132 family members are single-pass transmembrane proteins with a unique extracellular region containing a cohesin protein-protein interaction domain first described in prokaryotes (notably distinct from the similarly named eukaryotic regulator of mitosis, Cohesin), as well as bacterial immunoglobulin (BIG) repeats (Sanchez-Pulido and Ponting, 2018). The TMEM132 domain architecture is present across metazoans and choanoflagellates but not in plants or fungi, so it likely evolved in the lineage leading to animal multicellularity. Vertebrates have five TMEM132 orthologs, and functional analysis has identified roles for these factors in neural adhesion and cell migration. TMEM132A regulates collagen-binding integrin proteins and migratory behaviors in cultured cell types (Wang et al., 2021), and *tmem132A*-null mice fail to close the caudal neural tube, leading to spina bifida (Li et al., 2022). Additionally, multiple *tmem132* family genes are expressed in the developing central and peripheral nervous system in mice (Wang et al., 2022). Mutations in human TMEM132 family proteins have been implicated through genetic association studies to participate in a range of neurological conditions (Sanchez-Pulido and Ponting, 2018), including Anxiety Disorder (Erhardt et al., 2011; Erhardt et al., 2012; Haaker et al., 2014; Shimada-Sugimoto et al., 2016), hearing loss (Li et al., 2015; Liaqat et al., 2020), insomnia (Lane et al., 2017), and Major Depressive Disorder (Hu et al., 2021). Binding partners of TMEM132s and their downstream signaling targets have not been fully defined across systems, although TMEM132B can associate with the GABA(A) receptor to regulate alcohol responsiveness in mammals (Wang et al., 2024). Additionally, the sole *C. elegans TMEM132* ortholog regulates neuronal morphogenesis via the WAVE regulatory complex (Wang et al., 2021), though it remains unknown whether this mechanism accounts for all TMEM132 factors in other organisms. Our work therefore supports an ancient and enduring role for TMEM132 factors in regulating adhesion of neural cell lineages used for generating tissue connectivity.

Finally, the *tmem132-1(RNAi)* phenotype has implications in understanding the nature of homeostatic maintenance mechanisms in planarians. Because the *tmem132-1(RNAi)* phenotype does not appear to involve decay of detached OC cells or PRN aggregates, the identification of progenitors in the region between these two cell types suggests that each cell type might be capable of capturing and differentiating their own progenitors. In theory, signals from mature cells using homotypic adhesion to recruit their own progenitors could enable robust homeostasis and growth in complex environments that require the sorting of numerous distinct and spatially intermingled precursors for each lineage (Park et al., 2023).

Given the high diversity of progenitors and their broad spatial distributions in planarians, the problem of cell sorting into appropriate target locations is likely to be extensive and also crucial robust whole-body regeneration. Here, we have leveraged the planarian eye system to uncover mechanisms guiding *de novo* organ assembly in a post-embryonic context. Our analysis highlights two cell-surface factors, *ddr-1* and *tmem132-1*, likely operating on progenitors to enable interaction with the external environment, thereby directing eye progenitor targeting and assembly for robust eye regeneration.

## MATERIALS AND METHODS

### Experimental model

Asexually reproducing *Schmidtea mediterranea* (CIW4 strain) were cultured in 1x Montjuic salts at 18-20°C and fed pureed calf liver once a week. The animal colony was cleaned at least once every week for maintenance. Animals were starved for at least 7 days before experiments.

### Fluorescence-activated cell sorting (FACS)

Head blastemas from trunk fragments were surgically isolated from animals 5 or 8 days post-amputation. Animal fragments were rinsed in CMFB (CMF [400 mg/L NaH2PO4, 800 mg/L NaCl, 1200 mg/L KCl, 800 mg/L NaHCO3, 240 mg/L glucose,15 mM HEPES, pH 7.3] plus 1% BSA) supplemented with 0.01-0.05% NAC for up to 3 minutes, rinsed with CMF, rinsed with CMFB, and minced on an ice-cold petri dish with a fine scalpel. The suspension was transferred to an Eppendorf tube in CMFB supplemented with 1.0-2.0mg/ml collagenase and nutated for 20-30 minutes with vigorous pipetting every 5 minutes with a P1000. Finally, the solution was purified through a 40um mesh filter, centrifuged at 1250 rpm for 5 minutes, and resuspended in CMFB. For sorting, cell suspensions were incubated in Hoechst (10mg/mL, Invitrogen) and nutated in the dark for 45 minutes at room temperature. Prior to sorting, propidium iodide (3ul/mL, Sigma-Aldrich) was added to the suspensions. Propidium iodide negative cells were sorted based on DNA content using a FACSAria cell sorter. Cells from animals irradiated 3 days prior to sorting served as a negative control to determine gating parameters. X_1_, X_2_, and X-insensitive cells were collected from untreated animals and sorted into CMFB.

### Single-cell RNAseq library construction

FACS-sorted cells were pooled and manually counted using a C-Chip Hemocytometer (NanoEntek) on a Lecia DM5500B compound fluorescence microscope. Using the Chromium single cell 3’ and gel bead kit (10x Genomics), an estimated 25,000 cells were loaded into 2 lanes for the 8 dpa samples, and the chip was run on the Chromium X Controller (10x Genomics). For the 5 dpa samples, approximately 114,000 cells were loaded into 5 lanes using the Chromium HT single cell 3’ and gel bead kit, and the chip was run on the Chromium iX Controller. cDNA was generated and concentration was determined on a 4200 TapeStation System (Agilent) using a D5000 ScreenTape kit. Libraries from the cDNA samples were sequenced on Illumina NovaSeq X plus (Admera), and FASTQ files analyzed using the 10x Genomics Cloud Cell Ranger pipeline to align reads to the dd_v6 *S. mediterranea* transcriptome (Brandl et al., 2016), using Cell Ranger Aggr v9.0.1 to aggregate reads across samples. Loupe v9 software (10x Genomics) was utilized for data visualization and differential gene expression analysis based on sSeq (Yu et al., 2013).

### RNA interference (RNAi)

RNAi was performed by feeding dsRNA to animals every 2-3 days for the designated length of the experiment. Double stranded RNA (dsRNA) were produced by T7 RNA polymerase-mediated *in vitro* transcription reactions (NxGen, Lucigen) using PCR templates amplified from synthetic DNAs (Twist Bioscience). Control dsRNA was produced from a plasmid containing the *Caenorhabditis elegans unc-22* gene, a gene sequence not found in the planarian genome. Animals were fed liver mixtures containing 20% dsRNA, 5% food coloring dye, and 75% pureed calf liver.

### Fluorescence *in situ* hybridization (FISH) and immunostaining

FISH protocol used was performed as described in previous work (King & Newmark, 2013). Animals were fixed in 7.5% N-Acetyl-cysteine in PBS (w/v) followed by 4% formaldehyde in PBSTx (1X PBS+0.3% Triton X100, v/v), washed in PBSTx, and stored in methanol. Animals were bleached for 2-4 hours in a solution containing 5% deionized formamide (v/v) and 0.36% hydrogen peroxide (v/v) in 1xSSC. Digoxigenin-or fluorescein-labeled riboprobes were synthesized as described in previous work (King & Newmark, 2013) and detected with anti-digoxigenin-POD or anti-fluorescein-POD antibodies (1:2000, Roche/Sigma-Aldrich) blocked with 10% heat-inactivated Horse serum and 10% Western Blotting Blocking Reagent (Roche). Tyramide amplification was performed by depositing 0.2% rhodamine-tyramide or fluorescein-tyramide and 0.1% 4-iodophenylboronic acid (in dimethylformamide) in TSA buffer (2M NaCl, 0.1M Boric acid, pH 8.5). For double FISH, the enzymatic activity of tyramide reactions were quenched in 100mM sodium azide in TNTx. Nuclei were stained using 1:1000 Hoechst 33342 (Invitrogen) in TNTx. For planarian VC-1 ANTI-ARRESTIN immunostaining, animals were fixed in 4% formaldehyde as previously described (King & Newmark, 2013), blocked with 5% heat-inactivated Horse Serum in PBSTx, incubated in goat anti-mouse HRP (1:300) to detect labeling with mouse anti-ARRESTIN (1:10,000) before tyramide amplification.

### Image acquisition

Live animals were imaged using either a LeicaMZ125 dissecting microscope with a LeicaDFC295 camera or a LeicaS6D dissecting microscope with a FlexacamC3 camera. FISH-stained animals were imaged with a Stellaris 5 laser-scanning confocal microscope. Maximum projections were produced through the Leica software, and background signal was reduced as needed. Image adjustments including cropping, brightness, and contrast were made using FIJI/ImageJ.

### Quantification and statistical analysis of PCG expression

For analysis of PCG expression domains, maximum projection images were analyzed in ImageJ/Fiji to measure body length and length of the domain of PCG expression. For body length, animals were measured from the head tip to the end of the tail. PCG domain measurements were conducted by measuring the most anterior to most posterior expression for each gene. Statistical tests are indicated in figure legends and were conducted in Microsoft Excel.

### Mapping relative positions of eyes in live animals

Images of live or stained animals were overlaid on a grid, which normalized to the inter-eye distance such that the centers of left and right pigment cup cells were positioned at (-1,0) and (1,0) respectively. For analysis of ectopic eye phenotypes, the pigment cup of each ectopic eye was marked, and the distance from the original eyes was measured using WebPlotDigitizer (https://apps.automeris.io/wpd/) and then normalized in units equal to one half the inter-eye distance. The relative coordinates of each eye were extracted and plotted using ggplot2 in R.

## DATA AVAILABILITY

Single-cell transcriptome data was deposited to NCBI at PRJNA1503016.

## Acknowledgements

We thank members of the Petersen lab for critical comments and thoughtful discussion. We thank Prof. R. Zayas for the kind gift of the VC-1 antibody.

## Author Contributions

Chaeli Place: Investigation, methodology, validation, visualization, writing-original draft & preparation.

Katherine Lo: Investigation, methodology, validation, visualization, writing-original draft & preparation.

Eleanor Clark: Investigation, methodology, validation, visualization, writing-original draft & preparation.

Christian P. Petersen: Conceptualization, funding acquisition, methodology, resources, supervision, writing-original draft & preparation

Table S1. Single-cell RNAseq differential gene expression testing in eye cells

Loupe browser (v9) was used to conduct differential gene expression testing between the clusters of eye cells (eye neoblasts, eye post-mitotic progenitors, PRNs and OCs). Log2FC and padj values are indicated for each gene.

Table S2. Summary of FISH and RNAi screening of eye factors

Table summarizes genes tested for expression by double FISH to detect each gene with respect to a paired *ovo* riboprobe in day 5 regenerating trunk fragments as in Figure 1. Summary of notes and scoring for the FISH outcome is indicated. Summary of RNAi screening to detect live animal phenotypes and also animals fixed and stained with *opsin* and *ovo* riboprobes. Genes displaying enriched expression in eye progenitors, PRNs and/or OC cells based on the scRNAseq (padj<0.05 or log2FC>1) and also encoding receptors, membrane proteins, transcription factors, and signaling molecules were selected for the screening.

Table S3. DNA oligos used in the study

Genes tested in the screen were designed as synthetic DNA oligonucleotides with the indicated sequences.

---

**Figure S1.**
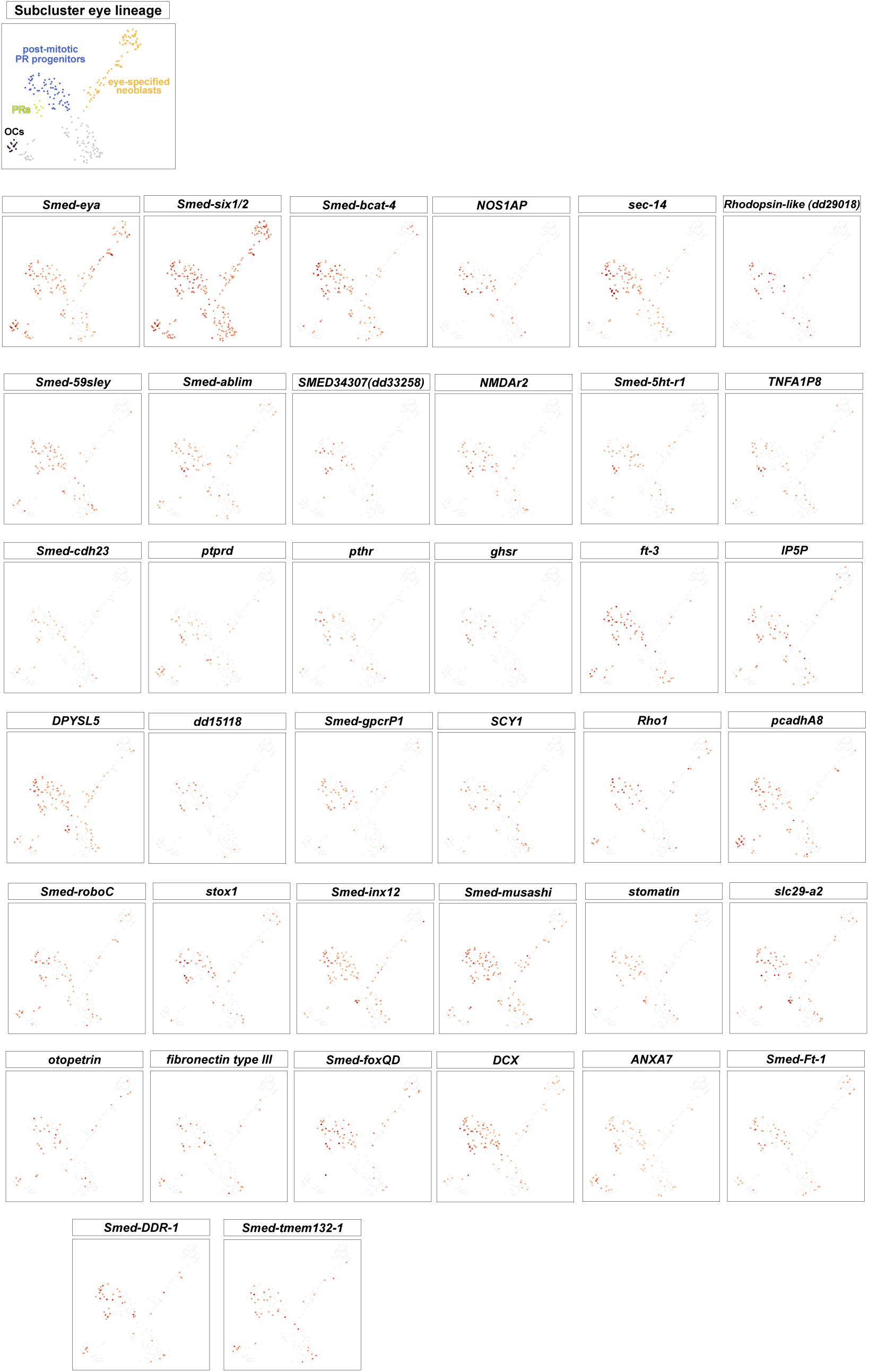
Expression of genes in eye cells by scRNAseq. Eye cells were subclustered by expression of *ovo* or *six-1/2* and displayed using 10x Genomics Loupe browser (v9), with cartoon showing assignment of clusters as in Figure 1 as *piwi-1*+ eye-specified neoblasts, *otxA+piwi-1*(low)*ovo*+ post-mitotic PR progenitors, differentiated PRs and differentiated OCs.

**Figure S2.**
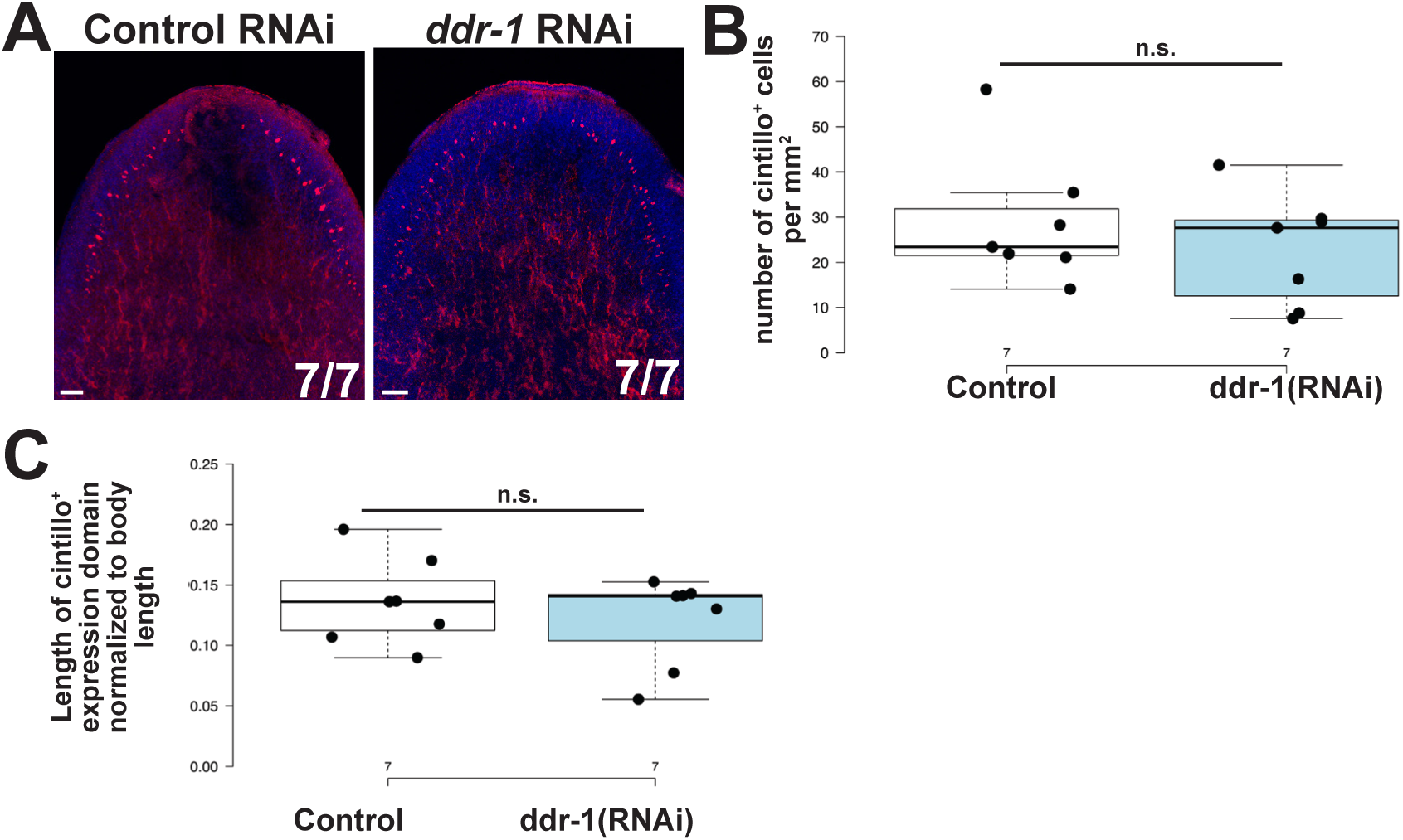
*ddr-1* RNAi does not affect head patterning. (A) *ddr-1* RNAi was conducted as in Figure 2, and animals stained for *cintillo* expression. (B-C) *ddr-1* RNAi did not increase or decrease the number of *cintillo+* cells normalized to body area (B) nor change the length of the *cintillo+* domain normalized to body length (C). Two-tailed unpaired t-tests were used for each comparison (n.s., p>0.05).

**Figure S3.**
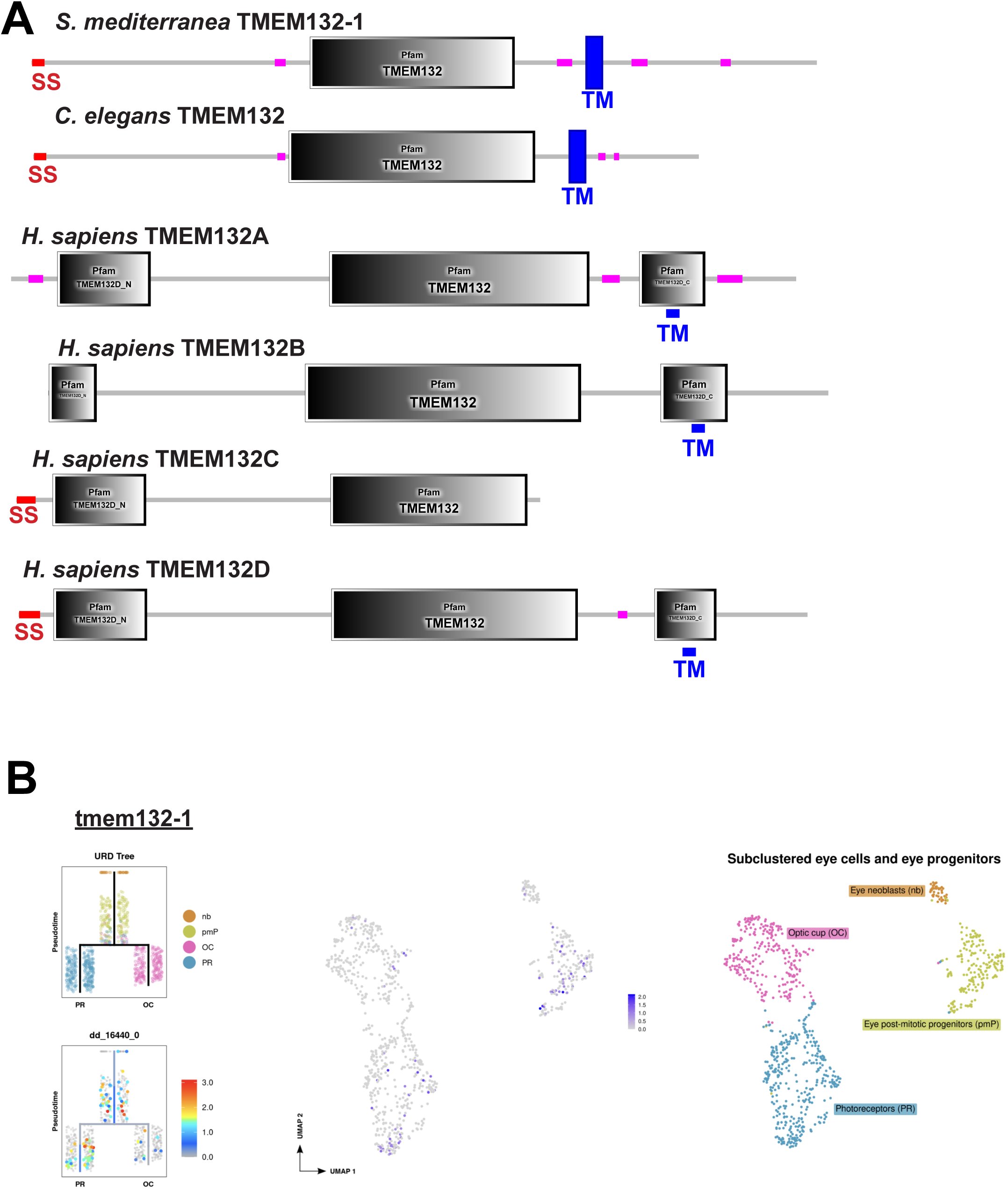
**TMEM132 domain structure and expression in eye cells**(A) Domain structures of TMEM132s from *C. elegans* and humans compared to planarian *tmem1321-1* using pfam/SMART (http://smart.embl-heidelberg.de/). SS, signal sequence; TM transmembrane domain. In human homologs, the TM domain is embedded in a TMEM132D_C domain. (B) Plots of *tmem132-1* (dd16440) as expressed in eye cells determined from a prior eye cell atlas (https://digieye.wi.mit.edu/) indicating clusters of eye neoblasts (nb), eye post-mitotic progenitors (pmP), differentiated photoreceptors (PR) and optic cups (OC). Left plot shows embedding of data with URD to predict a lineage relationship of each cell. Middle plot shows UMAPs from subclustered eye cells, and right cartoon shows assignment of cluster IDs in the middle panel. *tmem132-1* is expressed primarily in post-mitotic eye progenitors and PRNs.

